# Endocast and bony labyrinth of a stem gnathostome shed light on the earliest diversification of jawed vertebrates

**DOI:** 10.1101/2020.08.11.242974

**Authors:** You-an Zhu, Sam Giles, Gavin Young, Yuzhi Hu, Mohamad Bazzi, Per E. Ahlberg, Min Zhu, Jing Lu

## Abstract

Our understanding of the earliest evolution of jawed vertebrates depends on a credible phylogenetic assessment of the jawed stem gnathostomes collectively known as ‘placoderms’. However, their relationships, and even whether ‘placoderms’ represent a single radiation or a paraphyletic array, remain contentious. Here we describe the endocranial cavity and inner ear of *Brindabellaspis stensioi*, commonly recovered as a taxon of uncertain affinity branching near the base of ‘placoderms’. While some features of its braincase and endocast resemble those of jawless vertebrates, its inner ear displays a repertoire of crown gnathostome characters. Both parsimony and Bayesian analyses suggest that established hypotheses of ‘placoderm’ relationships are unstable, with newly-revealed anatomy pointing to a potentially radical revision of early gnathostome evolution. Our results call into question the appropriateness of fusiform ‘placoderms’ as models of primitive gnathostome anatomy and raise questions of homology relating to key cranial features.

**One Sentence Summary:** The skull of a 400-million-year old fossil fish suggests that hypotheses of early jawed vertebrate relationships might have to be turned on their head.

## Main text

One of the major transitions in vertebrate history was the evolution of gnathostomes, or jawed vertebrates, from jawless ancestors. The major morphological gap apparent when considering only living vertebrate diversity—extant jawless fishes comprise just hagfish and lamprey—is largely filled in by the fossil record (*1*). ‘Placoderms’, the most crownward assemblage on the gnathostome stem, occupy a pivotal place in this discussion. Traditional hypotheses of relationships posit a monophyletic Placodermi (*2–4*), whereas most recent analyses recover (*2–13*) ‘placoderms’ as a paraphyletic array from which crown gnathostomes arose (*5–11*); but see ref (*12*). In either scenario, taxa recovered near the base of the assemblage are typically dorsoventrally compressed, benthic forms—anatomically similar to agnathan outgroups—with fusiform, nektonic taxa recovered proximate to the gnathostome crown (*1,5,6,8–10*); but see ref (*11*). Uncertainty surrounding the relationships between different ‘placoderm’ groups, as well as their broader taxonomic status, are compounded by an uneven understanding of anatomy across the radiation, particularly of the phylogenetically informative braincase and brain cavity—endocast (see Supplementary Materials).

*Brindabellaspis stensioi* (*13*) is a ‘placoderm’-grade stem gnathostome from the Early Devonian of New South Wales, Australia. Although almost exclusively recovered among the earliest diverging ‘placoderms’ (*5,6–8,10–11,14–16*), it has variably been allied with rhenanids (*13*) acanthothoracids (*4*) and antiarchs (*12*), some of which are of dubious monophyly. Comparisons with jawless fishes have frequently been drawn on the basis of gross external and braincase anatomy (*13*) and general proportions of the endocast (*1, 17*). Other distinctive features, such as a large endolymphatic cavity, have been interpreted as autapomorphies (*13, 18*). Here, we provide high-resolution CT data of two more recently discovered specimens (Fig. 1, Fig. S1-4), detailing unexplored parts of the endocast and allowing previously described regions of the braincase and skull roof to be reinterpreted.

**Fig. 1.**
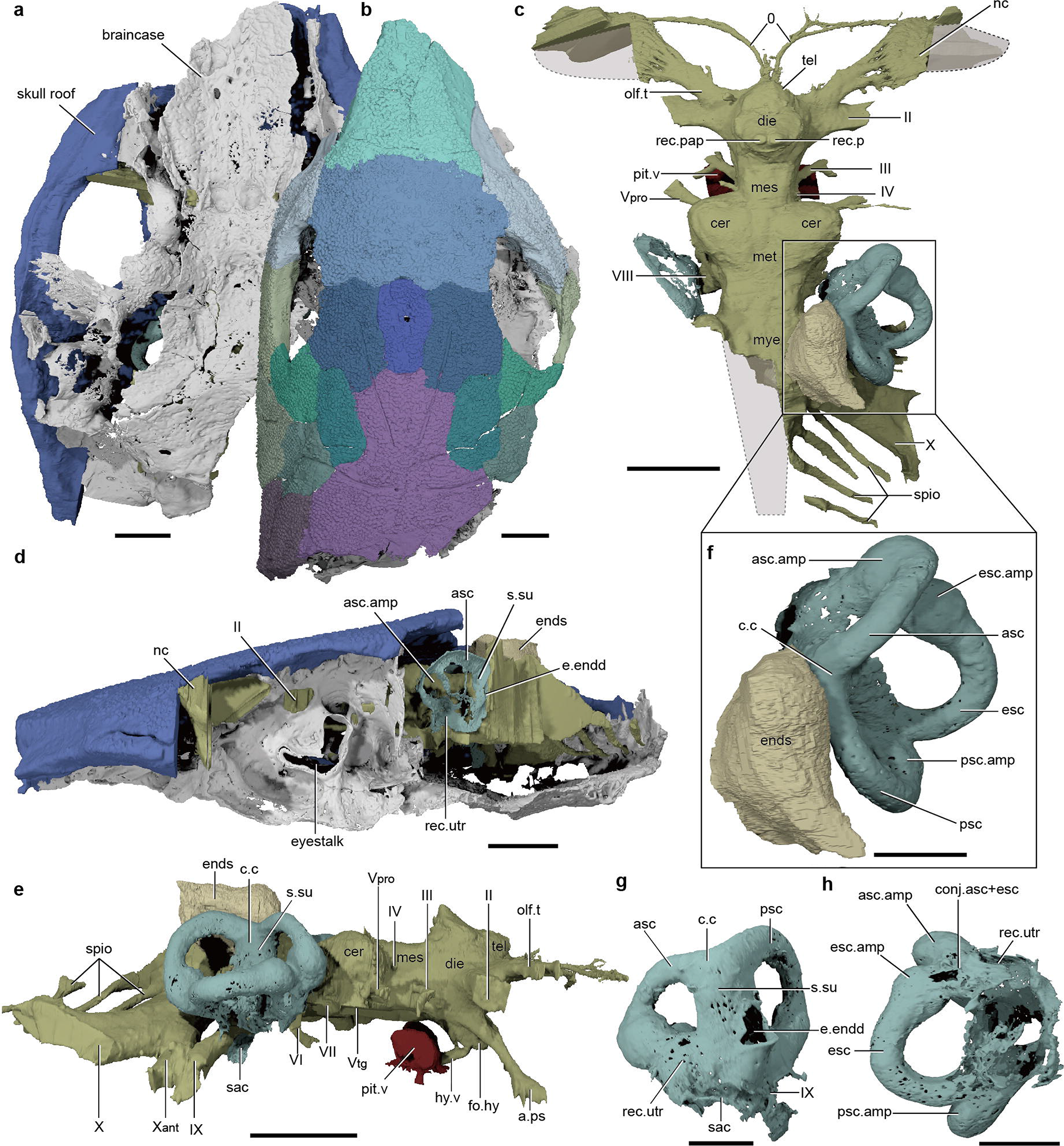
The skull of the ‘placoderm’ *Brindabellaspis stensioi*, based on high-resolution CT. **a**, Ventral view of endocranium (ANU 49493). **b**, Dorsal view of skull roof (AM F81911). **c,** Dorsal view of endocranial cavity (ANU 49493,); inset (**f**) shows bony labyrinth and endolymphatic sac region. **d**, Left lateral view of endocranium (ANU 49493). **e**, Right lateral view of endocranial cavity (ANU 49493). **g** and **h**, Mesial and ventral views of the skeletal labyrinth (ANU 49493). Abbreviations: a.ps, efferent pseudobranchial artery; asc, anterior semicircular canal; asc.amp, ampullae of anterior semicircular canal; c.c, crus commune; conj.asc+esc, conjunction of anterior and external semicircular canals; die, diencephalon; e.endd, exit of endolymphatic duct; ends, endolymphatic sac; esc, external semicircular canal; esc.amp, ampullae of external semicircular canal; fo.hy, hypophysial fossa; hy.v, hypophysial vein; mes, mesencephalon; met, metencephalon; mye, myelencephalon; nc, nasal capsule; olf.t, olfactory tract; pit.v, pituitary vein; psc, posterior semicircular canal; psc.amp, ampullae of posterior semicircular canal; rec.p, pineal recess; rec.pap, parapineal recess; rec.utr, utricular recess; sac, sacculus; spio, spino-occipital nerves; s.su, sinus superior; tel, telencephalon; 0, terminal nerve; II, optic nerve, III, oculomotor nerve; IV, trochlear nerves; Vpro, profundus branch of trigeminal nerve; Vtg, maxillary and mandibular branches of trigeminal nerve; VI, abducens nerve; VII, facial nerve; VIII, otic nerve; IX, glossopharyngeal nerve; X, vagus nerve; Xa, anterior branch of vagus nerve. Scale bars, **a-e**, 1 cm; **f-h**, 5 mm.

Tomographic data reveals the position of dermal bone sutures, clarifying the structure of the skull roof. Unlike in previous interpretations (*13,19*), we identify an independent median pineal plate sitting posterior to the rostropineal. We also confirm the presence of four bones contributing to the lateral margin of the skull roof, contra refs (*13,19*). *Brindabellaspis* possesses an elongate ossification (postmarginal) flanking the serial lateral line-bearing bones, resembling maxillate ‘placoderms’ (*6,10*) and early osteichthyans (*20*); in most other ‘placoderms’, the postmarginal is either much reduced or lost (*3*).

Broadly speaking, our results affirm past descriptions of the endocavity (*13*), although with key clarifications and additions. The extremely short telencephalic region of the endocast has a flat anterior face with no bulge anterior to the olfactory and terminal nerves, contra ref (*13*) (Fig. 1c and e). CT data also clarify that, as in crown gnathostomes (*8,21,22*), the ventral sides of the telencephalic and diencephalic cavities are smooth and continuous between the optic nerve and hypophysis, rather than displaying a step-like transition as in most ‘placoderms’ (*18,23*) (Fig. S5). We also confirm that the hypophysis is oriented anteriorly (*13*) (Fig. 1e). Although an anteriorly (or ventrally) directed hypophysis has sometimes been considered restricted to agnathans and ‘placoderms’(*18*), it is also reported in crown gnathostomes (*16,22,24*). Characters previously suggested as being shared between agnathans and *Brindabellaspis*, such as a laterally expansive cerebellum and anteroposteriorly elongate vagus complex (Fig. 1d and f, Fig. S3), are now known to be widespread in other stem and crown gnathostomes (*16,18,22*) (Fig. 2 and Fig. S5), and are presumably plesiomorphic for the gnathostome crown. As described by Young (13), the olfactory tracts are elongate, and diverge anteriorly towards the widely separated and laterally positioned nasal capsules (Fig. 1c and Fig. 2, Fig. S3a-d). Divergent olfactory tracts are otherwise only known in crown gnathostomes (*16,21,22*) in other ‘placoderms’ and the galeaspid *Shuyu* (*25*) the olfactory tracts are parallel and typically short (Fig. 2). The myelencephalic region of the endocast anterior to the vagus nerve, which is usually proportionately long in most ‘placoderms’ (Fig. 2 and Fig. S5d-g) but short in agnathans and crown gnathostomes (Fig. 2 and Fig. S5a-c, h-m), appears intermediate in length in *Brindabellaspis*.

**Fig. 2.**
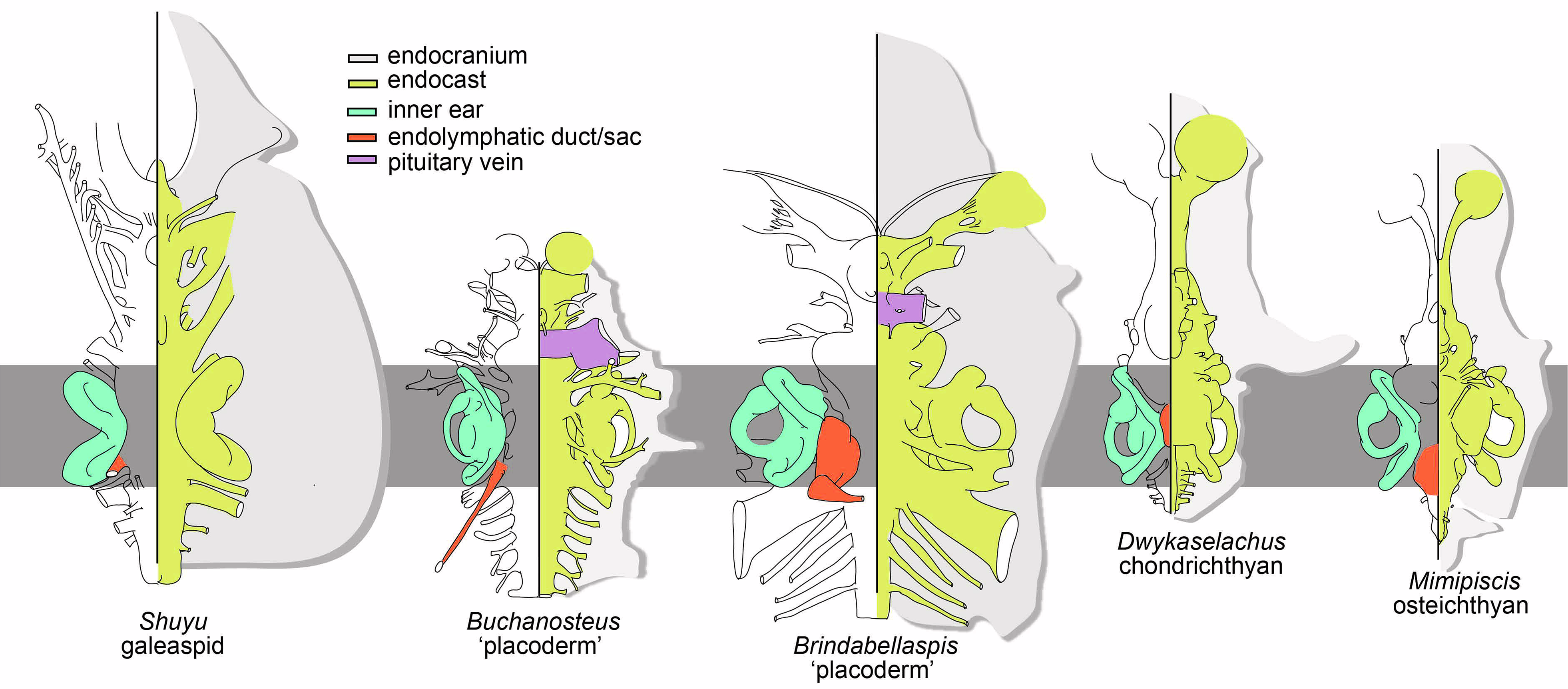
Comparative morphology of crania in selected early vertebrates, aligned and scaled to match skeletal labyrinth dimensions (grey bar). Cranial outlines (light grey) and endocast outlines (yellow) in dorsal (left) and ventral (right) views. Data sources for each genus are provided in Supplementary Information.

Our CT data reveal important new anatomical details of the bony labyrinth and endolymphatic complex. In addition to features identified in the endocast, the bony labyrinth of *Brindabellaspis* (Fig. 1c-f, Fig. 2, Fig. S4) bears unexpected similarities to those of crown gnathostomes, with considerable difference to those of other ‘placoderms’. The labyrinth is anteroposteriorly short, and all three semicircular canals have large diameters. The anterior semicircular canal is significantly shorter than its posterior counterpart, and in dorsal view the two diverge at a much smaller angle than in other ‘placoderms’ such as the rhenanid *Jagorina* and arthrodire *Kujdanowiaspis* (Figs. 1c-f and 3, Fig. S4). Most strikingly, CT data demonstrate that the anterior and posterior semicircular canals of *Brindabellaspis* join in a crus commune, with a pronounced sinus superior developed ventrally. This configuration is typical of crown gnathostomes (*16,21,22*), and the combination is unknown in other ‘placoderms’ (Fig. 3). There is no significant preampullary portion of the posterior semicircular canal, and the utriculus does not separate the anterior and external semicircular canals (both contra the condition in all known ‘placoderms’ except *Romundina* (*18*)). While incomplete ventrally, the curvature of the sacculus suggests that it is significantly smaller than in other ‘placoderms’, barely protruding laterally (Fig. 2 and Fig. 3). It is also restricted ventral to the plane of the external semicircular canal, a condition seen elsewhere only in crown gnathostomes (Fig. 2 and Fig. 3). Despite the lack of an external semicircular canal or utricular chamber in agnathans (*25*), a number of labyrinth characters can be polarised across the jawless-jawed vertebrate transition. Osteostracans possess small angles between anterior and posterior semicircular canals, and a crus commune but no developed sinus superior (Fig. 3).

**Fig. 3.**
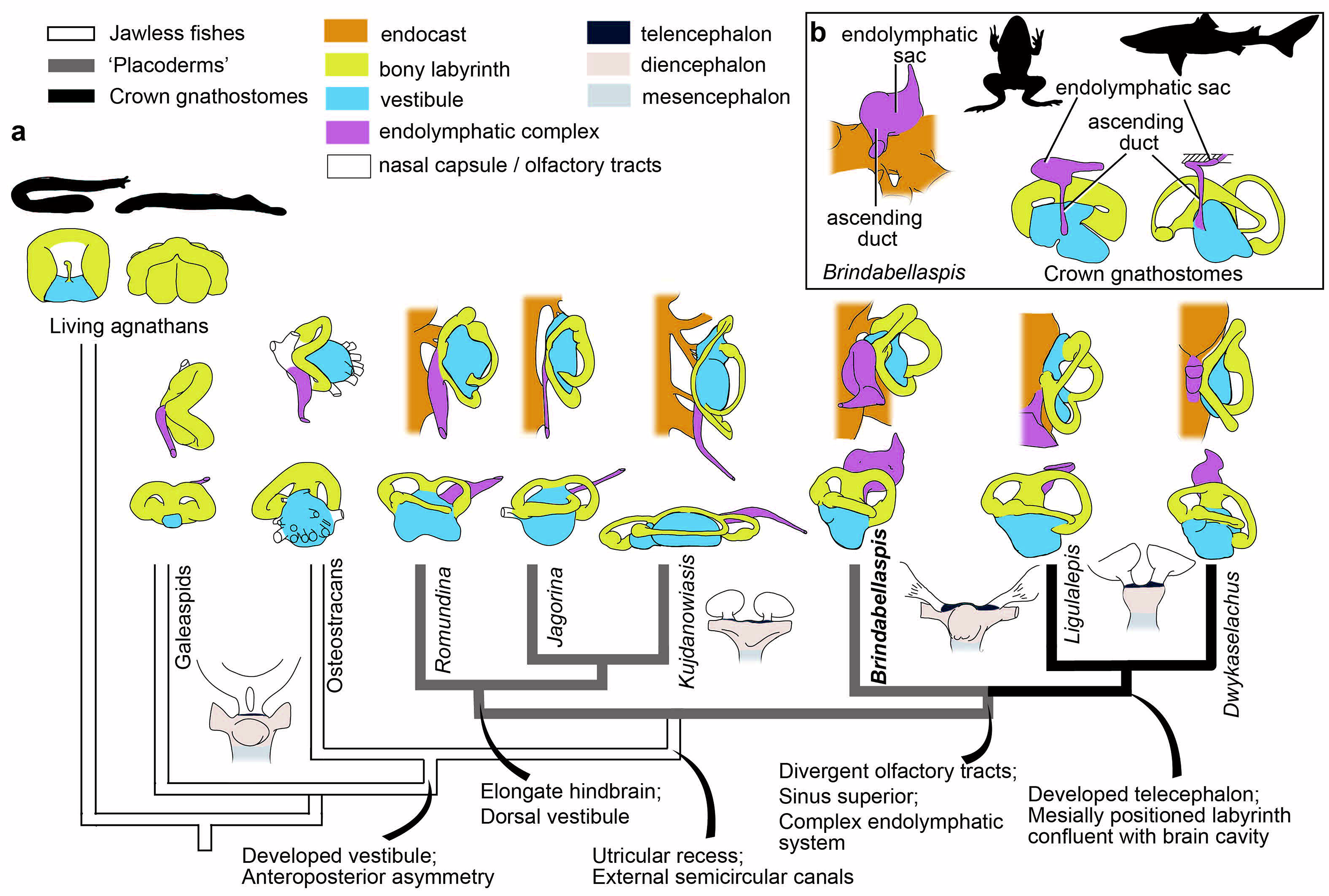
Summary phylogeny showing the evolution of the inner ear and endolymphatic complex in early vertebrates. **a**, Simplified phylogeny from the parsimony strict consensus tree (Fig. S6a). **b**, The endolymphatic complex of *Brindabellaspis* and selected crown gnathostomes in lateral view, showing the shared characters including an ascending duct and an endolymphatic sac.

Although previously considered autapomorphic (*13*), our data allow similarities to be drawn between the endolymphatic systems of *Brindabellaspis* and crown gnathostomes. The complex in *Brindabellaspis* can be divided into three distinct sections: a large, well-developed endolymphatic sac; an ascending duct connecting the vestibular chamber to the endolymphatic sac; and a distal duct extending from the sac, through the dermal bone, and opening externally (Figs. 1d-f, Fig. 2 and Fig. 3, Fig. S4a, d-f). There is no “second sac”, contra ref (*18*). In both jawless and jawed stem gnathostomes, the endolymphatic complex is a simple tube-like structure that extends unidirectionally, and is positioned close to the labyrinth (*17,18,23,25*). In contrast, the crown gnathostome system is more complex and divided into three distinct regions (*26*), much as in *Brindabellaspis*, and located mesially, closer to the brain cavity than the labyrinth (*27,28*) (Fig. 2 and Fig. 3).

A revised and expanded morphological matrix, analysed under both parsimony and Bayesian frameworks, provides novel—and conflicting—insights into early gnathostome evolution. Under parsimony analysis, jaw-bearing gnathostomes fall into one of two monophyletic groups (Fig. 3 and Fig. S7a). The more stemward of these contains the bulk of traditionally-recognised ‘placoderms’, albeit with arthodires representing a nested radiation within this clade.

Unexpectedly, *Brindabellaspis* is recovered as the earliest diverging member of a clade comprising, successively: antiarchs, maxillate ‘placoderms’, and crown gnathostomes. The position of antiarchs as proximate to the gnathostome crown, with arthrodires representing an independently fusiform radiation, is unexpected (*1,5,6,8–10*); but see ref (*12*), and perhaps indicates the importance of endocranial data and previous biases towards external morphology. The endocavities of *Minicrania* (*29*) and *Phymolepis* (*30*) hint at the presence of a mesially-directed endolymphatic duct and an endolymphatic sac, as well as a relatively short hindbrain. Although not included in the phylogenetic analysis, these anatomical similarities between the endocrania of antiarchs and *Brindabellaspis*—and, by extension, the gnathostome crown—lend support to the hypothesis of relationships suggested in our parsimony results. However, support values amongst early gnathostomes are low, and the proximity of antiarchs to the gnathostome crown node raises several questions of homology. The transition from posteriorly-positioned to anteriorly-positioned nasal capsules, as well as changes to jaw suspension, are now optimised as evolving twice: once within the clade comprising *Romundina*, rhenanids, ptyctodonts, petalichthyids and arthrodires; and once within the clade comprising maxillate ‘placoderms’ and crown gnathostomes. The recovery of arthrodires as removed from maxillate ‘placoderms’ plus crown gnathostomes also requires a number of homoplasies in the skull roof and trunk armour.

Results under Bayesian analyses differ from our parsimony analysis and recall more common hypotheses of placoderm paraphyly (*1,5,6,8–10*), with antiarchs recovered as the earliest-diverging ‘placoderm’ clade and arthrodires as sister taxa to maxillate ‘placoderms’ and the gnathostome crown. Outside of these nodes, however, other ‘placoderms’—including *Brindabellaspis*—fall in a polytomy, and arthrodires are recovered as paraphyletic. Resolving this conflict represents a fundamental challenge of early gnathostome evolution, and is one that cannot be resolved without detailed CT-based reassessment of the anatomy of key ‘placoderm’ taxa.

Our work adds considerably to knowledge of labyrinth and endocast variation across stem gnathostomes, highlighting the major impact that CT-based descriptions and re-examination of key taxa can have on both phylogenetic resolution and schemes of morphological evolution. The unexpected character combination in *Brindabellaspis* suggests that endocranial characters previously considered exclusive to crown gnathostome are likely widely distributed amongst a diversity of stem jawed vertebrates. However, outstanding questions remain about the homology of features common to both arthrodires and crown gnathostomes, notably in the skull roof and nasal capsules. The conflicting phylogenetic hypotheses of relationships presented here highlight major uncertainties on the gnathostome stem, calling into question long-standing assumptions about patterns of character evolution. Recent work on the diversity of ‘acanthothoracid’ dentitions, also revealed by CT data, suggests a more complex picture of dental character evolution and provides independent evidence that at least some ‘acanthothoracids’ may branch closer to the gnathostome crown node than previously thought (*11*). Notably, the position of arthrodires as removed from maxillate ‘placoderms’ plus crown gnathostomes challenges previous installations of fusiform, arthrodire-like taxa as a representative of the primitive gnathostome condition (*10,23,31*).

## Acknowledgements

We thank M. Turner and T. Senden for CT scanning support, and M. Brazeau for discussion.

## Funding

This work was supported by the Strategic Priority Research Program of Chinese Academy of Sciences (XDB26000000, XDA19050102), the National Natural Science Foundation of China (41872023, 41530102) and Australian Research Council Discovery grants (DP0772138, DP1092870). Y.Z., P.E.A. and M.B. were supported by Swedish Research Council grant 2014-4102 and a Wallenberg Scholarship from the Knut and Alice Wallenberg Foundation. S.G. was supported by a Royal Society Dorothy Hodgkin Research Fellowship.

## Author contributions

G.Y., J.L. and Y.Z. designed the research project; J.L., S.G., Y.Z. and Y.H. carried out the reconstruction of datasets and digital segmentation; Y.Z., S.G. and J. L. performed the phylogenetic analysis and constructed figures; M.B. and S.G. performed principal component analyses; Y.Z., J.L., S.G., and M.B. wrote the manuscript; and all authors reviewed and revised the manuscript.

## Competing interests

The authors declare no competing interests.

## Data availability

The CT data that support the findings of this study, as well as 3D surface files of described material, are shared privately in figshare for review: https://figshare.com/s/d040d38b2e0ae3501f65. All other data files are included in the Supplementary Materials.

## Supplementary Materials

*Materials and Methods*

*Supplementary Text*

*Figures S1-S6*

*Captions for Data S1 to S5*

## Notes

### Competing Interest Statement

The authors have declared no competing interest.

