## Supplementary material for "Endocast and bony labyrinth of a stem gnathostome shed light on the earliest diversification of jawed vertebrates": SI text

**Affiliations:**
[1]Key Laboratory of Vertebrate Evolution and Human Origins of Chinese Academy of Sciences, Institute of Vertebrate Paleontology and Paleoanthropology, Chinese Academy of Sciences, Beijing 100044, China.

[2]Subdepartment of Evolution and Development, Department of Organismal Biology, Uppsala University, Norbyvägen 18A, SE-752 36 Uppsala, Sweden.

[3]School of Geography Earth and Environmental Sciences, University of Birmingham, Birmingham, UK, B15 2TT.

[4]Department of Earth Sciences, Natural History Museum, Cromwell Road, London SW7 5BD, UK

[5]Department of Applied Mathematics, Research School of Physics, Australian National University, Canberra, ACT, Australia 2601.

[6]Australian Museum Research Institute, Sydney, NSW, Australia 2010.

[7]CAS Center for Excellence in Life and Paleoenvironment, Beijing 100044, China.

[8]University of Chinese Academy of Sciences Beijing 100039, China.

**This PDF file includes:**

Materials and Methods
Supplementary Text
Figs. S1 to S6
Captions for Data S1 to S5
References

### Materials and Methods

Material

Young assigned six specimens to *Brindabellaspis stensioi*, three of which are skulls (*13*). Five additional specimens, which also provide information on skull morphology, were described in 2018 (*19*). The endocranial morphology of *Brindabellaspis* redescribed here is based mainly on X-ray computed microtomography of two specimens: AM F81911 and ANU 49493. We list below the six original and five new specimens. Additional 'CPC' numbers, used by Young (*13*) for some ANU specimens, are given in parentheses for completeness. All but the last two specimens are held in the Department of Applied Mathematics, Research School of Physics, ANU.

ANU 37 (CPC 16991), left AL and scapulocoracoid; ANU 48 (CPC 16990), incomplete SM plate; ANU 1677 (CPC 16987), incomplete skull with some perichondral bone attached (Holotype); ANU 1678 (CPC 16988), eroded skull revealing much of the endocranial cavity; ANU 1679 (CPC 16989), posterolateral corner of a skull; ANU 2584, incomplete fractured and distorted skull that shows a complete posterior margin; ANU 3247, slightly distorted anterior portion of the skull roof with underlying perichondral ossifications, broken off posteriorly at the anterior edge of the orbit and nasal cavity; ANU 1224, incomplete fractured anterior portion of the skull roof; ANU 49493, completely acid-etched partial skull and braincase; AM F81911 (Australian Museum, Sydney), partial skull and braincase, partly acid-etched; NHMUK PV P50287 (Natural History Museum, London), left IL plate.

X-ray computed microtomography

Both specimens were scanned with the HeliScan CT Scanner (ANU1) in the CT Lab, Department of Applied Mathematics, Research School of Physics, Australian National University. ANU 49493 was scanned in 2017 with a 3 mm aluminium filter (resolution: 30 microns). The specimen was placed 161 mm from the source, with detector position 625 mm from the source, and was probed separately with a polychromatic X-ray beam (Bremsstrahlung radiation) with accelerating voltage of 120kV and a current of 120$\mu$A. AM F81911 was first scanned in 2017 with a 3 mm aluminium filter (resolution: 34 microns), with the specimen placed 180 mm from the 120kv accelerating voltage and 120 $\mu$A current polychromatic X-ray beam source; the detector was placed 786 mm from the source. AM F81911 was re-scanned in 2018 to obtain a more focused view of the inner ear area, using the same filter as the 2017 scan (resolution: 35 microns). The specimen was positioned 187 mm from the source, and detector position positioned 599 mm from the source; it was probed separately with 100kv accelerating voltage and 150$\mu$A current polychromatic X-ray beam (Bremsstrahlung radiation). All reconstructions of these scans were based on 3600 radiographic projections on a 2872 × 2840 Pixium Flat Panel camera.

Phylogenetic analysis

Parsimony analysis was performed in PAUP v. 4.0a158 (*32*) with the following settings: 500 random addition sequences, five trees held at each step, maxtrees set to automatically increase, nchuck =10 000, chuckscore =1, tree bisection and reconstruction strategy enabled using a data matrix with a total of 355 characters and 116 taxa. The data matrix is based on refs 8,10 and 33. We added or amended 14 characters (both new and from ref. 34), revised a number of codings and removed redundant characters to give a total of 116 taxa and 360 characters. The four taxa in Vaškaninová et al. (*11*) were not included due to the low proportion of characters that can be coded on the basis of published data; all four act as wildcard taxa. Osteostraci and Galeaspida were set as the outgroup. All characters are unweighted, and six characters (c.62, c.123, c.160, c.256, c.258, c.262) were ordered. The full character list is given below. Bremer support values were calculated in TNT, with only Bremer Indices higher than one retained in the strict consensus tree nodes. An optimization tree, showing all ambiguous character changes, was generated in MacClade (*35*) (see Supplementary Data 5). The Bayesian analysis was run in MrBayes v.3.2.6 (*36*) under the Mkv model, with Galeaspida set as the outgroup. The analysis was run until the standard deviation of split frequencies reached less than 0.01, indicating convergence had been reached, and this was confirmed in Tracer (*37*). The first half of each run was discarded as burn-in.

The results of both the parsimony and Bayesian analyses are summarized in Supplementary Figure 7. The parsimony analysis resulted in 39601 most parsimonious trees of 1102 steps. The complete strict consensus tree with node support is summarised in Supplementary Data Figure 6a. RI = 0.791; CI = 0.352; HI=0.648; RC=0.278. The Bayesian analysis is summarized in Supplementary Data Figure. 6b.

### Supplementary Text

Anatomical notes on new skull roof interpretation

The original identification of skull roof bones in *Brindabellaspis* (*13*) has been revised twice (*3,19*) on the basis of external examination. Tomographic data reveals newly recognized sutures, allowing a revised interpretation of the skull roof pattern. This differs significantly from previous reconstructions in the following details:

Firstly, previous interpretations have reconstructed a large rostropineal plate with a posterior convex margin, resembling the plate in the acanthothoracid *Romundina* and arthrodires such as *Buchanosteus*. In our new interpretation, however, this posterior extension is in fact a separate pineal plate. The rostral has a sinuous posterior margin, which interdigitates with the narrow, rectangular pineal and the large, quadrate preorbital (Fig. S2a).

Secondly, previous interpretations do not agree on the bones around the posterolateral boundary of the orbit and the lateral edge of the skull roof. Our new interpretation resembles that of Young (*3*), with a series of four bones (contra the five in refs. *13,19*) forming the lateral edge of the skull roof: a postnasal; a bar-shaped bone forming the ventral margin of the orbit and tentatively identified as a postorbital plate; a postmarginal; and a paranuchal (Fig. S2a). The postmarginal canal runs through the postmarginal, rather than through the presumed postorbital plate (as reconstructed by refs. *13,19*).

The skull roof of *Brindabellaspis* is highly apomorphic, particularly with regard to the anteriorly elongated bill-like premedian and accompanying lateral postnasals. Interestingly, its skull roof pattern shows some resemblance to the recently discovered Silurian maxillate placoderms, *Entelognathus* and *Qilinyu* (*6,10*). In both *Brindabellaspis* and maxillate placoderms, a large postmarginal is present and forms a substantial portion of the lateral margin of the skull roof, outflanking the serial bones bearing the lateral line canal (carried by the postorbital, marginal and anterior and posterior paranuchals). In most other placoderms, the postmarginal is either much reduced, forming only the tip of the lateral corner of the skull roof, as in many arthrodires, or completely lost, as in

*Romundina*, petalichthyids and ptyctodonts. Notably, some of the earliest-diverging osteichthyans such as *Guiyu* primitively have large ossifications flanking the bones bearing the lateral line canals. These bones form a considerable part of posterolateral margin of the skull roof (*20*), although the exact homology of these two bones (named extratemporal and accessory extratemporal in *Guiyu*) to the postmarginal of placoderms is difficult to establish. Finally, in *Brindabellaspis* the lateral-line-bearing marginal and anterior paranuchals are a series of rounded bones like in *Entelognathus* and *Qilinyu*, rather than the large polygon or irregularly shaped bones in most other placoderms.

Notes on endocast and labyrinth morphology in stem gnathostomes

The endocranial cavity morphology seen in 'placoderms' highlights the extreme diversity present across the assemblage, and with reference to outgroups allows us to reconstruct character polarity.

The two groups of jawless stem gnathostomes (or "ostracoderms") that have well-preserved endocranial cavities, namely osteostracans and galeaspids, display considerable variation in morphology. The close affinity of osteostracans to jawed vertebrates is supported by features of the inner ear, in particular its drum-shaped vestibular cavity (sacculus), and anteroposterior asymmetry (*38-40*). In contrast, the galeaspid *Shuyu*, despite possessing separated nasal sacs and olfactory tracts as in jawed vertebrates, lacks vestibular structures; the ventral part of the labyrinth, or *pars inferior*, is simply a continuation of the anterior and posterior semicircular canals as in living cyclostomes (*25*). The whole profile of the inner ear in *Shuyu* also resembles living cyclostomes in being anteroposteriorly symmetrical. Both *Shuyu* and osteostracans possess a distinct crus commune, but the crus commune of osteostracans does not appear to extend into a developed sinus superior. The ventral extension of the crus commune seems variable in the osteostracans with inner ear morphology known. It is somewhat more pronounced in *Kiaeraspis auchenaspidoides* (Fig. 3; ref. *38*:text-fig. 19) than in *Mimetaspis hoeli* (ref. *38*:text-fig. 18) and *Norselaspis glacialis* (ref. *40*: fig. 30), but is still more similar to the condition in *Romundina* than the distinct sinus superior in *Brindabellaspis* and crown gnathostomes. A better understanding of the crus commune in osteostracan inner ears is hampered by the lack of updated digital data, the availability of which would allow dissection of individual structures in strictly directional views.

Arthrodires represent some of the first described and "stereotypical" placoderms (*41-45*). In recent phylogenetic analyses, arthrodires are typically resolved close to the crown gnathostome node (*46*-*50*), but in our parsimony analysis are nested within a monophyletic group containing many conventional 'placoderms'. The endocranial cavity, especially the labyrinth morphology, is well-known in *Kujdanowiaspis* (*44*, *51*, *52*) and *Buchanosteus* (*53*), both of which display derived (or apomorphic) features when compared to agnathan outgroups. These features include a very long hindbrain; elongated general profile of the labyrinth; absence of crus commune, and laterally directed endolymphatic duct (Fig. 3). In agnathans and most 'placoderms' the endolymphatic duct runs parallel in dorsal view (Fig. 2 and Fig. 3). The telencephalic and diencephalic regions of the arthrodire endocast resemble *Romundina* in having a bulged dorsal "forehead" and a ventral pre-hypophysial "step" anterior to the hypophysial fossa (Fig. S5). The olfactory tracts proximally join the robust and long optic nerve canals that are perpendicular to the axis of the endocast. The condition in which the olfactory tracts proximally join the robust and long optic nerve canal is shared in *Shuyu*, other 'placoderms' such as *Jagorina* (*54*), and *Brindabellaspis* (Figs. 1d, 2 and 3, Fig. S3), suggesting it is the primitive gnathostome condition. Of special interest is the incompletely known endocast in the derived arthrodire *Tapinosteus* (44). *Tapinosteus* belongs to the arthrodire subgroup eubrachythoraci (*55*) and supposedly adapted to free-swimming lifestyle with fusiform body, resembling crown gnathostomes. However, the preserved part of the inner ear resembles that of other arthrodires in the low general profile and the absence of crus commune. The olfactory tracts are also parallel. Although data are limited, this indicated that endocast and inner ear morphology are unlikely to broadly reflect ecological mode.

Together with arthrodires, antiarchs were among the first 'placoderms' to be described (*41*). Antiarchs are typically resolved at the base of the 'placoderm' position close to *Brindabellaspis* in recent phylogenetic analyses. Notably, antiarchs possess various features difficult to compare to other 'placoderm' subgroups, such as a unique skull roofing pattern and "lateral plate", a cheek bone bearing the dental and occlusal surface, a pectoral appendage clad in macromeric dermal plates, and a long, box-like trunk armour with multiple median dorsal plates. It is unclear whether these features represent derived or primitive gnathostome conditions—or indeed apomorphies—partly due to the lack of neurocranial evidence in the group. Endocavity anatomy is known in most detail for *Minicrania* (*56*), a primitive antiarch from the Early Devonian of Yunnan, China. External observation of the specimen suggests the presence of a bulbous endolymphatic sac not unlike the one in *Brindabellaspis*. The putative endocavity in *Minicrania* shows a short hindbrain section, and, based on the imprint of the braincase on the visceral surface of the skull roof, it is likely that most antiarchs display this condition. The distal part of the endolymphatic duct in the yunnanolepiform antiarch *Phymolepis* (*57*) is mesially directed as in *Brindabellaspis*, providing potential further support for a close relationship between antiarchs, *Brindabellaspis* and the gnathstome crown. In addition, antiarchs appear to have an anteroposteriorly compact labyrinth, with the anterior and posterior semicircular canals in dorsal view forming an angle close to that in *Brindabellaspis* and most crown gnathostomes, and smaller than that in petalichthyids and arthrodires. It is worth noting that these observations are not included in the current data matrix due to the limited data available for *Minicrania* and *Phymolepis*. Detailed descriptions based on tomographic data of this and other taxa is likely to yield more support for the systematic position of antiarchs, providing more insights into the primitive gnathostome characters in the future.

Petalichthyids include morphologically diverse taxa such as *Diandongpetalichthys*, *Eurycaraspis* and macropetalichthyids, but are traditionally united in one group. To date, the endocranial cavity is only reported in the macropetalichthyids *Macropetalichthys* and *Shearsbyaspis*. The former is more completely preserved (*52*). *Shearsbyaspis* is digitally reconstructed based on tomographic data, but the posterior part of cranium in the examined specimen is lost (*58*). The olfactory tracts in both taxa are unusually long, but are anteriorly directed and parallel, as in most other 'placoderms', and the endocast in *Macropetalichthys* also displays an elongate hindbrain region contributed by both the otic and the cranio-spinal sections. In both taxa, the large and bulbous sacculus, thin semicircular canals and pre-ampullary portions of the anterior and external semicircular canals resemble those of arthrodires and rhenanids (*58*). *Macropetalichthys* is unique in having the distal part of the endolymphatic duct vertically ascends and flares. This cavity was labelled “fosse endolymphatique” by Stensiö (*52*), following chondrichthyan terminology. However, unlike the chondrichthyan condition, the endolymphatic duct first extends posteriorly for most of its length, and this flared section is far more distal than the endolymphatic sac in *Brindabellaspis*, resulting in unclear homology. However, the three-dimensional morphology of endolymphatic complex in *Macropetalichthys* is poorly known, and more information is needed to determine whether the flared condition is independent from those in *Brindabellaspis* and crown gnathostomes or not.

Rhenanids are ‘placoderm fishes’ with flat, disc-shaped bodies like those of stingray, and the dermal skeleton deviates considerably from the typical ‘placoderm’ bauplan. As in *Macropetalichthys*, the endocranial cavity of the rhenanid *Jagorina* lacks an updated three-dimensional description, despite good preservation of the neurocranium (*52*, *54*, *59*). The inner ear of *Jagorina* resembles arthrodires in displaying a bulbous utricular and sacculus chamber, anterior and external semicircular canals separated by the utricular recess, and a simple tube-like endolymphatic duct. The olfactory tracts are coded as “diverged” in current matrix based on drawings by Stensiö (*54*), the otic section of the hindbrain is long as in most other ‘placoderms’, and the craniospinal section is very short as in crown gnathostomes.

*Brindabellaspis* was previously assigned into Acanthothoraci, a poorly known and loosely defined placoderm group showing considerable morphological disparity, and which is almost certainly paraphyletic. The labyrinth of the acanthothoracid *Romundina* displays an interesting combination of inner ear and endolymphatic features (*60*), representing a possible intermediate between *Brindabellaspis* and arthrodire / rhenanid conditions. The general profile is not elongated, but is not as longitudinally compressed as in *Brindabellaspis* and the crown group. The angle between the anterior and posterior semicircular canals in dorsal view is sharper than the condition in rhenanids and arthodires, but is larger than that in agnathans, crown-group, and *Brindabellaspis*. The crus commune is present but the sinus superior is not developed, more-or-less resembling the condition in osteostracans. The posterior semicircular canal does not have an extensive preampullary section; the sacculus is large, dorsally positioned in relation to the external semicircular plane, and is flat inclined in anterior view; and the auditory nerve bifurcates before entering the labyrinth cavity, all as in other placoderms. However, the sacculus is irregularly shaped, and the utricular does not separate the ampullary ends of anterior and external semicircular canals, which joins before entering the utricular chamber, as in *Brindabellaspis* and crown gnathostomes (Figs. 2 and 3).

Notes on homoplasies invoked by our parsimony analysis

Although supported by our parsimony analysis, the phylogenetic distance between arthrodires and maxillate ‘placoderms’ plus the gnathostome crown demands a number of homoplasies. These broadly fall into three categories: jaw suspension (c.92, c.93, c.328), pectoral fin articulation (c.194), and the posterior nose – anterior nose transition. In previous hypotheses of relationships, the posterior nose to anterior nose transition (an anatomical complex of several characters: c.112, c.115, c.127, c.134 c.157) happened only once: at the node subtending arthodires, maxillate ‘placoderms’ and crown gnathostomes. In the scenario of relationships hypothesised in our parsimony analysis, early members of each of the two jawed vertebrate clades are ‘posterior nosed’, with the transition to an anterior nose occurring independently within them (once in maxillate ‘placoderm’ and crown gnathostomes, and in a stepwise manner in the clade comprising Romundina, rhenanids, ptyctodonts, petalichthyids and arthrodires). There are also three homoplasies between the skull roof and trunk armour of arthrodires and maxillate ‘placoderms’ (c.92, c.313, c.323, c.328).

**Extended Figure Captions.**

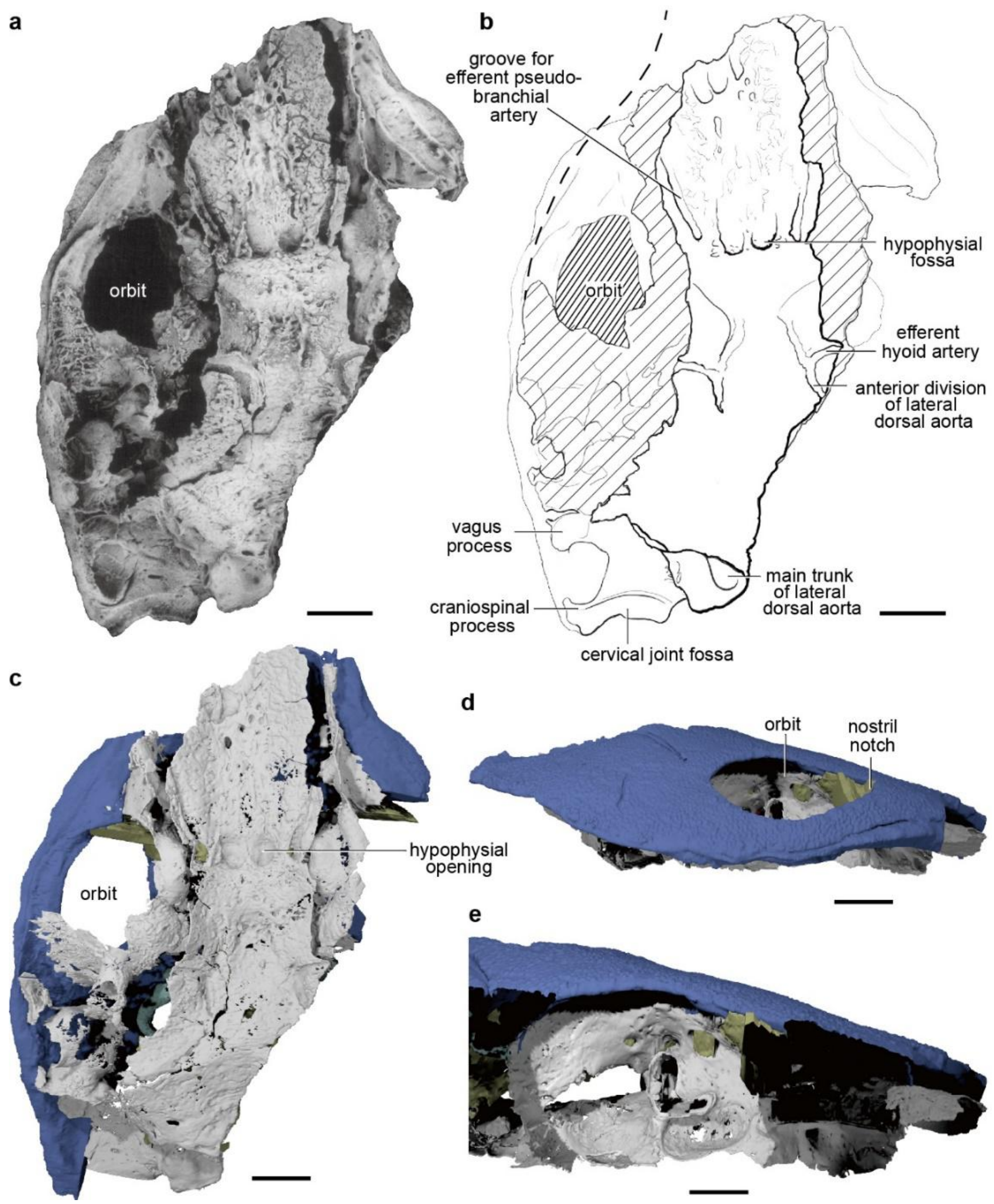


**Fig S.1** Braincase of *Brindabellaspis stensioi* (ANU 49493). **a-b**, Ventral view (photo, **a**) and interpretative drawing (**b**). **c-e**, Segmented braincase based on high-resolution CT in ventral (**c**) and right lateral view (**d**), with dermal bone cropped away in order to expose the orbital region (**e**). Scale bar, 1 cm.

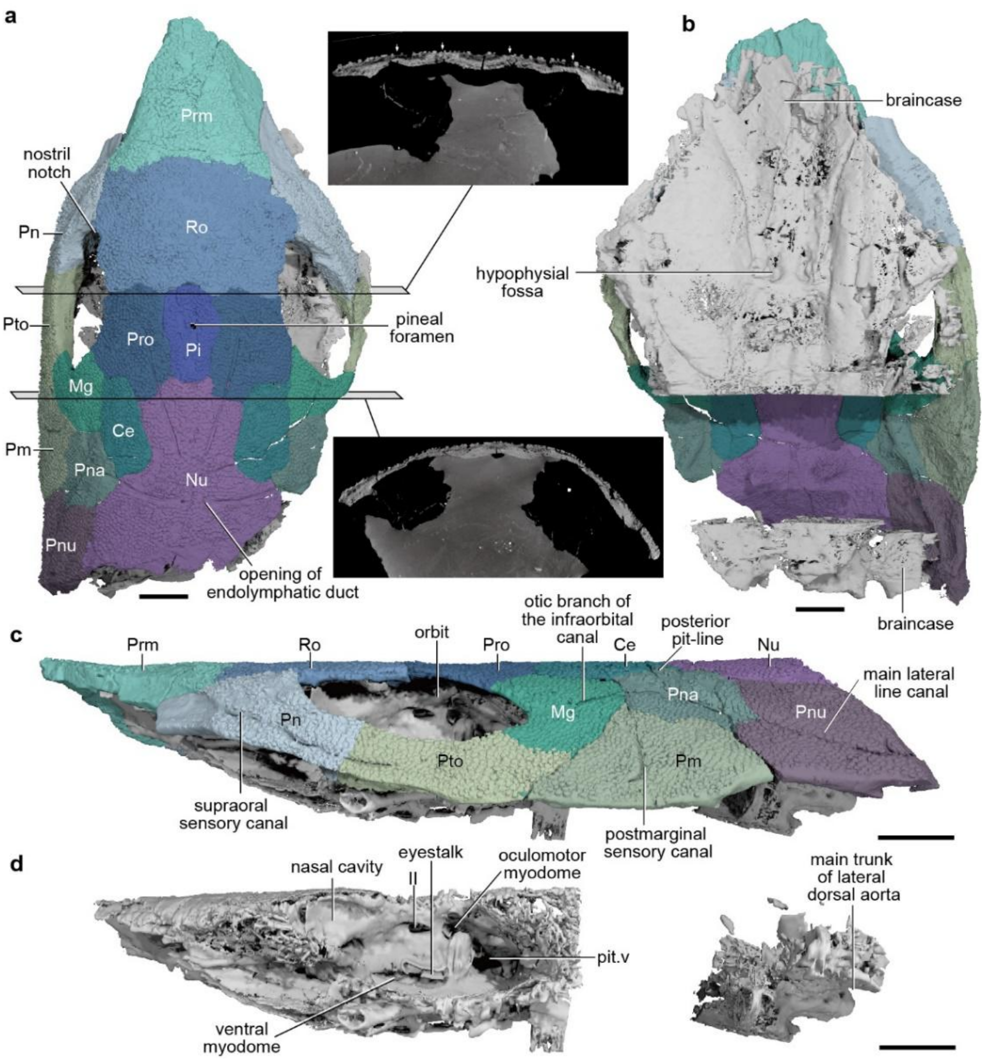


**Fig S.2** Braincase of *Brindabellaspis stensioi* (AM F81911), based on high-resolution CT. **a**, Dorsal view, showing the dermal skull roof. **b**, Ventral view, showing the segmented skull roof and neurocranium. **c**, Left lateral view of whole skull. **d**, Left lateral view of neurocranium only. Segmenting could not be completed in the otic region of the braincase due to very low contrast between the braincase and surrounded limestone matrix, which was only partially acid-etched. Digital transverse sections showing the dermal bone sutures. Abbreviations: Ce, central; Mg, marginal; Nu, nuchal; Pi, pineal; Pm, postmarginal; Pn, postnasal; Pna, anterior paranuchal; Pnu, paranuchal; Prm, premedian; Pro, preorbital; Pto, postorbital; Ro, rostral. Scale bar, 1 cm.

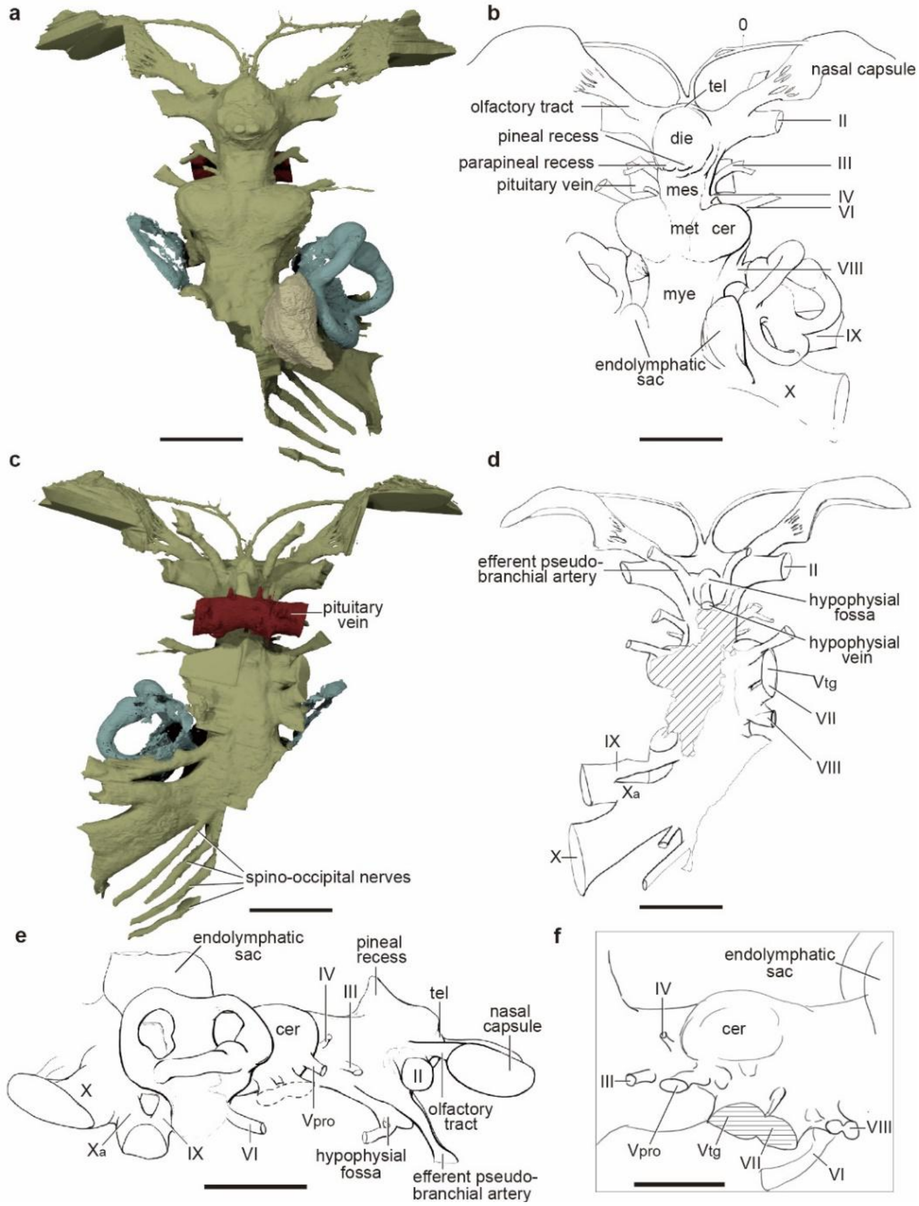


**Fig S.3** Endocast and associated nerve canals of *Brindabellaspis stensioi* (ANU 49493) based on high-resolution CT. **a**, Rendering of endocast in dorsal view. **b**, Interpretative drawing of endocast in dorsal view. **c**, Rendering of endocast in ventral view. **d**, Interpretative drawing of endocast in ventral view. **e**, Interpretative drawing of endocast in right lateral view, and showing the details of the trigeminal complex (**f**). Abbreviations see Figure 1. Scale bars, **a-e**, 1 cm; **f**, 5 mm.

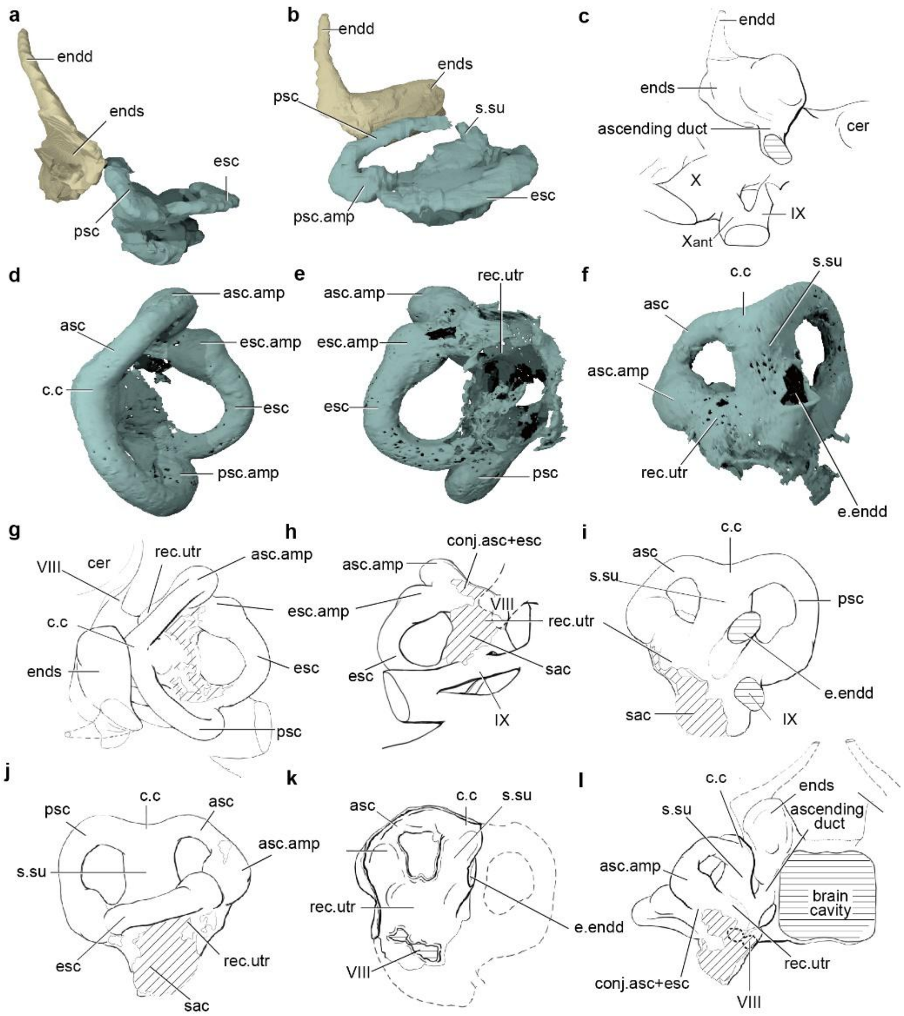


**Fig S.4** Inner ear and endolymphatic complex in *Brindabellaspis*. **a-b,** Rendering of the right skeletal labyrinth and endolymphatic system (AM F81911) in posterior (**a**) and lateral (**b**) views, showing the posterodorsal connection of the endolymphatic sac to the distal tube. **c-l,** Rendering and interpretative drawing of the skeletal labyrinth (ANU 49493). Interpretative drawing of right skeletal labyrinth in right lateral view with inner ear removed (**c**). Right skeletal labyrinth in dorsal (**d**, rendering; **g**, with endolymphatic complex, interpretative drawing), ventral (**e**, rendering; **h**, interpretative drawing), mesial (**f**, rendering; **i**, interpretative drawing), lateral (**j**, interpretative drawing), and anterior (**l**, with endocast and endolymphatic complex, interpretative drawing) views. **k**, Interpretative drawing of left skeletal labyrinth in lateral view. Abbreviations: endd, endolymphatic duct. Other abbreviations see Figure 1. Not to scale.

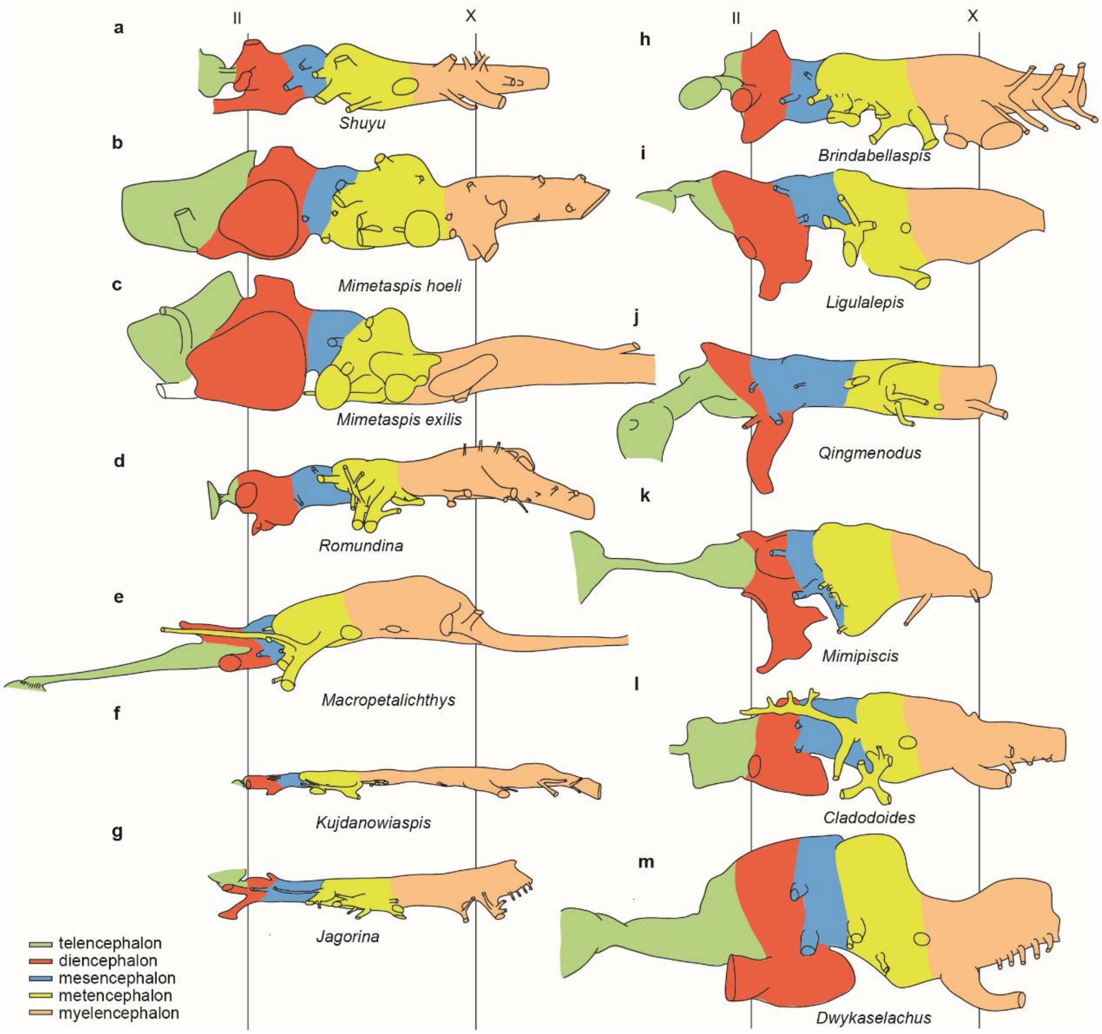


**Fig S.5** Comparative morphology of the endocast in selected early vertebrates, showing the proportions of different regions in lateral view. The uncoloured line drawings sketch the ventral contour of the telencephalic and diencephalic cavity. In taxa without an otic wall separating the brain cavity and labyrinth, the boundary between the metencephalon and myelencephalon is estimated by the position of the utricular recess, where the cranial nerve VIII connects to the labyrinth. **a-c**, Jawless fishes. **d-h**, 'placoderms'. **i-k**, Osteichthyans. **l-m**, Chondrichthyans. Not to scale. Anterior to the left.

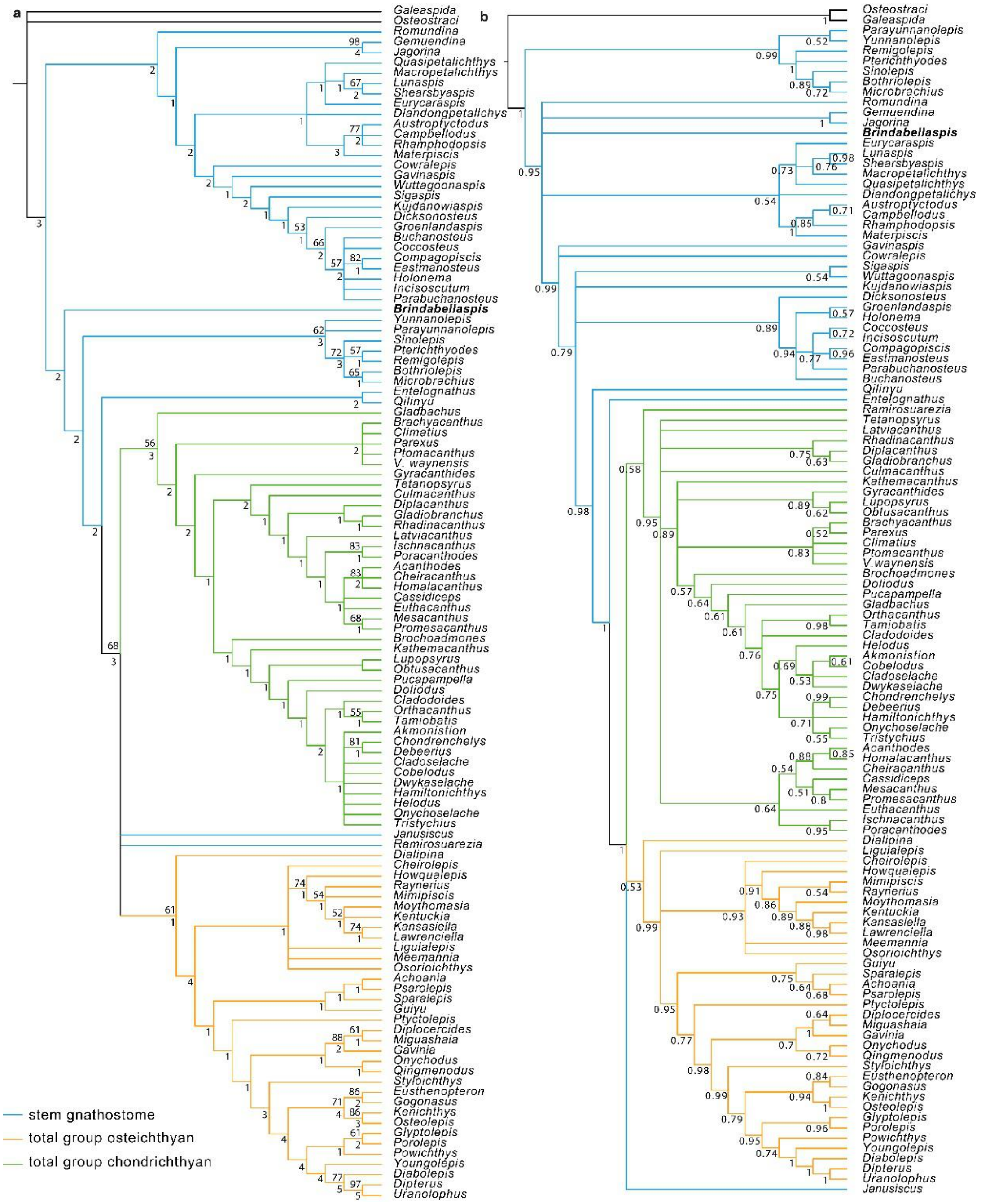


**Fig S.6** Consensus trees from phylogenetic analyses. **a**, Strict consensus of 39601 most parsimonious trees (1102 steps) for 116 taxa and 360 equally weighted characters. Digits above nodes indicate Bremer decay indices above 1. **b**, Maximum clade credibility tree from the Bayesian analysis of the same dataset. Values associate with nodes indicate the posterior probabilities.

**Data S1. (separate file)**

Phylogenetic analysis: list of taxa and character list.

**Data S2. (separate file)**

Nexus file

**Data S3. (separate file)**

MrBayes file

**Data S4. (separate file)**

Blender file containing labyrinths of selected early gnathostomes

**Data S5. (separate file)**

Single most parsimonious tree showing all character optimisations.
